# *In vitro* floral development in poplar: Insights into seed trichome and trimonoecy regulation

**DOI:** 10.1101/2022.10.15.512362

**Authors:** María A. Ortega, Ran Zhou, Margot S.S. Chen, William P. Bewg, Bindu Simon, Chung-Jui Tsai

## Abstract

Woody perennials including *Populus* spp. (poplars) have a juvenile phase that ranges from several years to decades in length. This and the year-long floral development process are major impediments to breeding and to fundamental research of reproductive traits. Here we report a CRISPR-empowered *in vitro* flowering system and demonstrate its application using three reproductive traits: sex, seed trichomes, and a previously undescribed potential for trimonoecy in poplar.

## Main text

Several regulators of floral development, such as LEAFY (LFY) and FLOWERING LOCUS-T (FT), have been used to induce precocious flowering in annual models and fruit trees (Weigel & Nilsson, 1995; Callahan *et al*., 2016). Translating these findings into the poplar system has met with various challenges, including dwarfism and sterility (Zhang *et al*., 2010). Using heat-inducible promoters for *FT* expression has circumvented many of the developmental anomalies; however, repeated application of heat treatments over weeks or months can be detrimental to microsporogenesis (Hoenicka *et al*., 2014). The method’s efficacy is also season- and genotype-dependent, limiting its widespread adoption (Zhang *et al*., 2010; Hoenicka *et al*., 2016).

Here we present an induction-free, *in vitro* flowering system by targeting a negative regulator of floral initiation, *CENTRORADIALIS* (*CEN*, also called *TERMINAL FLOWER*), for knockout (KO). CEN antagonizes FT and LFY to regulate meristem determinacy and flowering (Bradley *et al*., 1997; Jaeger *et al*., 2013). RNAi-silencing of *CEN* orthologs in poplar and pear (*Pyrus communis*) shortened their juvenile phase to under three years (Mohamed *et al*., 2010; Freiman *et al*., 2012), and CRISPR-KO of *CENs* in kiwi (*Actinidia chinensis*) reduced flowering time to under one year (Varkonyi-Gasic *et al*., 2019). For the present work, we adopted CRISPR/Cas9 to edit the *CEN1*/*CEN2* paralogs in a female *Populus tremula* × *alba* INRA 717-1B4 hybrid (hereafter 717). *In vitro* flowering of rooted plantlets was observed under long-day tissue culture conditions within four months of *Agrobacterium* transformation. During vegetative propagation of the mutants, single flowers developed directly from axillary buds of subcultured stem segments and decapitated mother plants within 1-2 wk (Fig. 1a). Amplicon-sequencing confirmed all 17 flowering events as *cen1cen2* double-KOs (Data. S1). *In vitro* flowering was reproducible in nodal cultures where carpels with a cupular disk developed directly from axillary buds (Fig. 1b). This effectively fast-tracked multi-season floral organogenesis to a timeframe of days. We next asked whether both paralogs are involved in floral development as only *CEN1* transcripts are detected in shoot tissues (Fig. S1). We generated 10 *cen1* single-KO events with biallelic mutations (Data. S1) and observed *in vitro* flowering phenotypes similar to the *cen1cen2* double-KOs (Fig. S2). The findings suggest that CEN1, but not CEN2, represses poplar floral development, as independently reported by another group (Sheng *et al*., 2022).

**Figure 1.**
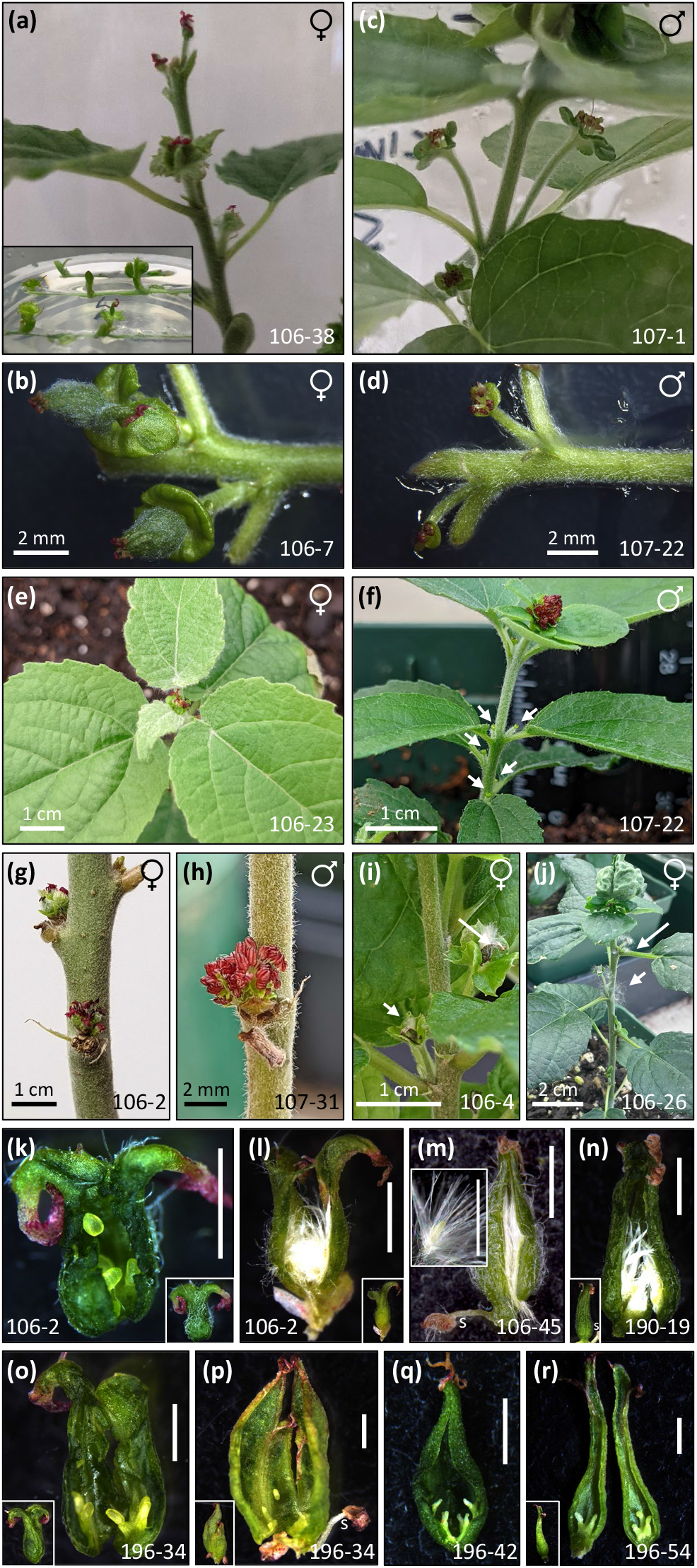
Rapid flowering of poplar mutants. (a, b) *In vitro* female flowers of *cen1cen2* mutants from a decapitated plant (a), subcultured stems (a inset), or node cultures (b). (c, d) *In vitro* male flowers of *arr17cen1cen2* mutants from a decapitated plant (c) or nodes (d). (e-h) Female (e, g) and male (f, h) flowers in terminal (e-f) and axillary (f-h) buds following soil transplanting (e-f) or cutback (g-h). (I, j) Mature seedpod-like capsules with cottony trichomes (i) and their release (j). (k-n) Bisected *in vitro* carpels of *cen1cen2* (k-m) or *cen1* (n). Intra-ovarian trichomes were not observed in immature carpels (k) but abundant in later stages and remained attached to ovules causing their comose appearance (m inset). (o-r) Bisected *in vitro* carpels of *cen1cen1myb* glabrous mutants devoid of intra-ovarian trichomes. Insets are pre-bisected carpels. Independent events are labeled. Bars in (k)-(r), 1 mm. s, stamen.

A female-specific, type-A response regulator (*ARR17*) gene was recently identified as the sole sex determinant in poplars with either the XY (most *Populus* spp.) or ZW (*P. alba*) system (Müller *et al*., 2020). *ARR17*-KO in an early-flowering female *P. tremula* triggered male flower development (Müller *et al*., 2020). Interestingly, 717 has a hybrid sex configuration (♀, XZ) derived from *P. tremula* (♀, XX) and *P. alba* (♂, ZZ), which is supported by detection of hemizygous *ARR17* in the haplotype-resolved 717 draft genome (Phytozome). Monoallelic *ARR17*-KO is predicted to convert 717 from female to male, and this was tested using multiplex-editing of *ARR17* and *CENs* to determine the sex-switch outcome *in vitro*. We observed male flowers in all eight *arr17cen1cen2* events (Fig. 1c-d) and confirmed targeted mutations in all cases (Data. S1). Our results support *ARR17-*dependent sex switch in a XZ background. The fast-track flowering system is more time- and labor-efficient than the inducible-*FT* method (Zhang *et al*., 2010; Hoenicka *et al*., 2016; Müller *et al*., 2020) for studying floral traits in poplars.

Soil-transplanted female and male mutants (8-10 events per group) flowered in terminal as well as axillary buds after acclimation (Fig. 1e-f). The indeterminate-to-determinate meristem conversion resulted in cessation of vegetative growth, accelerated maturation with thickening and darkening of preexisting leaves, and following repeated cutbacks, prolific root suckers not seen in WT (Fig. S3). Healthy suckers and sprouts grew vegetatively for several weeks before they terminated into flowers, whereas cutback triggered flowering from axillary buds within days (Fig. 1g-h). Unfertilized carpels still matured into seedpod-like capsules, which eventually opened and released cottony hairs (Fig. 1i-j). These phenotypes remained consistent across all mutant lines, over multiple rounds of transplanting, and for over a year.

To further exemplify utility of *in vitro* flowering in reproductive trait investigation, we asked whether a group of MYB transcription factors recently shown to be essential for leaf and stem trichome initiation (Bewg *et al*., 2022) also regulate seed trichome development. Seeds with tufted hairs are characteristic of poplars and willows, with the cottony trichomes facilitating wind dispersal of seeds. In urban and plantation forestry, seed hairs are carriers of airborne allergens representing a potential health hazard (Hu *et al*., 2008). Multiplex-KO of *CENs* and trichome-regulating *MYBs* produced 11 early-flowering (♀) glabrous events (Fig. S4), with confirmed edits at all 12 (eight *MYB* and four *CEN*) target sites (Data. S1). We compared *in vitro* carpel development between trichome-bearing and trichomeless (♀) mutants. Ovules were already visible in the immature carpels of *cen1cen2* we bisected (Fig. 1k). Intra-ovarian trichomes were not observed until carpels reached *c*. 2 mm in length and gradually filled the ovary during maturation (Fig. 1l-n). An abundance of trichomes remained attached to ovules which resembled comose seeds (Fig. 1m inset). In glabrous mutants, ovules but not intra-ovarian trichomes were observed throughout carpel development in all three events examined (Fig. 1o-r). The results suggest seed trichomes, like other aerial organ trichomes, are regulated by the same MYBs in poplar, and provide a molecular basis for engineering hairless seeds for genetic confinement or for reducing allergen spread in urban/plantation forestry.

Finally, monitoring of *cen1cen2* nodal cultures revealed a remarkable diversity of sex morphs in 717 (♀, XZ), with male, female, and perfect flower development (trimonoecy) dictated by stem node position on the mother plant. In multiple *cen1cen2* events, we observed male flowers with stamens only from subapical nodes (Fig. 2a-c), perfect flowers with carpels and stamens from upper nodes (Fig. 2d-f, see also Fig. 1m,n,p), female flowers only from middle nodes (Fig. 2g-i), and vegetative buds from older nodes (Fig. 2j-l). The unusual stamen appearance (♂ or ⚥) was transient, limited to just a few nodes near the top. Such a developmental gradient is difficult to capture in soil-grown *cen1cen2* mutants because of the pleiotropic phenotypes discussed above. Nevertheless, carpellate flowers with stamens or stamen-carpel chimeras were occasionally observed and always near the top of the (cutback) plant (Fig. S5), consistent with transient male organ development. The occasional appearance of perfect flowers has also been reported in the field (Boes & Strauss, 1994; Zhang *et al*., 2010). Overall, the data suggest a critical role for ontogenic regulation on poplar sex determination that warrants further research. In sum, the *in vitro* flowering system bypasses the multi-year reproductive phase transition in poplar and fast-tracks the year-long floral development process to days and weeks. It offers a facile model for investigating floral traits and holds promise for rapid-cycle breeding and genomic selection in perennial trees.

**Figure 2.**
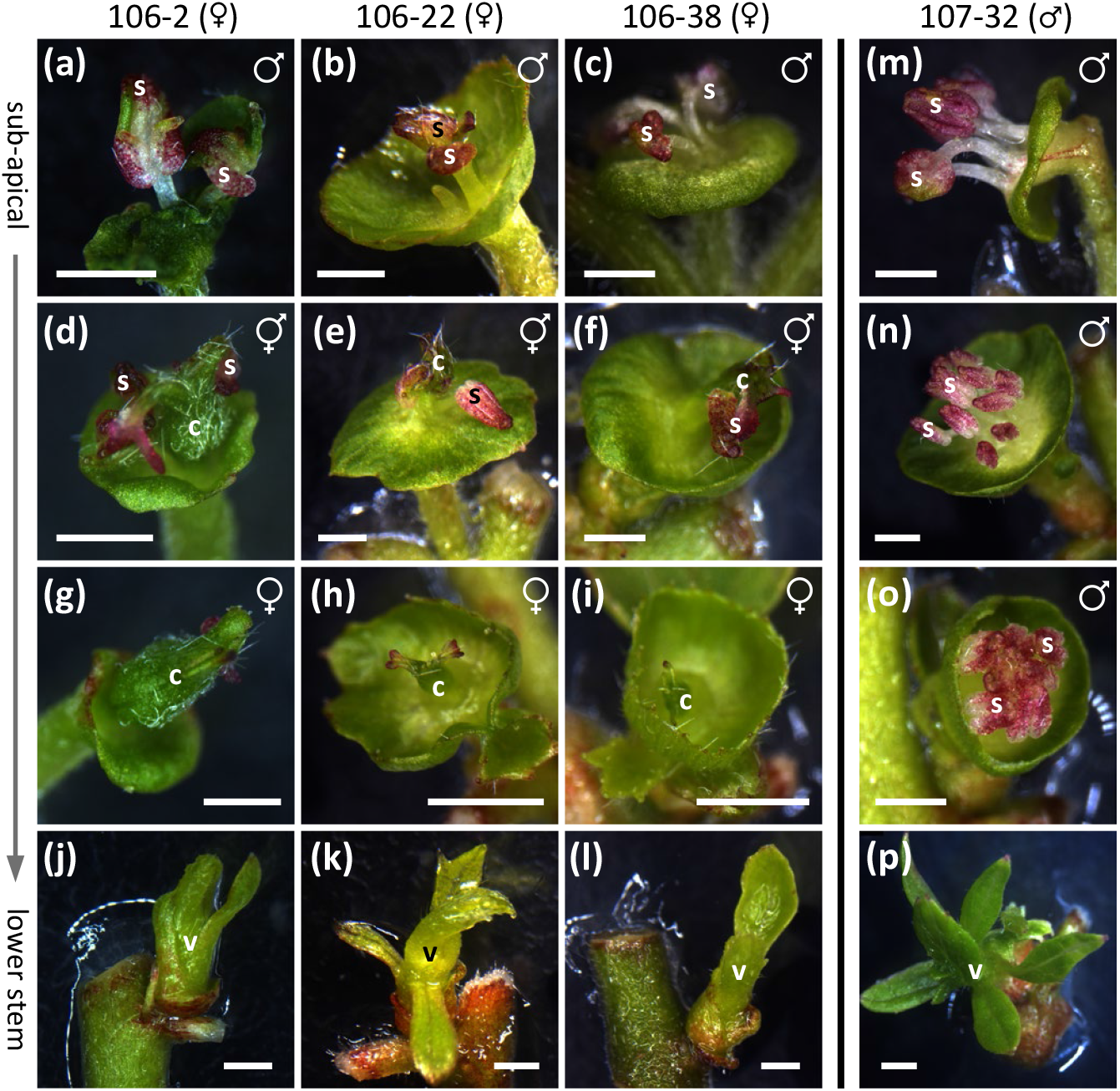
Ontogenic regulation of sex morphs in *cen1cen2* (♀) mutants. (a-c) Male flowers with stamens (s) from subapical nodes. (d-f) Perfect flowers with both stamens and carpels (c) from upper nodes. (g-i) Female flowers from middle nodes. (j-l) Vegetative (v) buds from lower nodes. Three independent events are shown. (m-p) Corresponding nodes from an *arr17cen1cen2* (♂) mutant for reference. All images were acquired 5-7 d after node culture except (p) at 18 d. Bars, 1 mm.

## Acknowledgments

We thank Gilles Pilate of INRAE, France for providing poplar clone 717, and Makenzie Drowns and Brent Lieb for amplicon library assistance.

## Author contributions

C.-J.T. and R.Z. conceived the study, C.-J.T. and M.A.O. designed the experiments, M.S.S.C., M.O.A., W.P.B., R.Z. and B.S. conducted the experiments, C.-J.T. wrote the paper with contributions from M.A.O. and R.Z.

## Supporting Information

Table S1. gRNA, oligo, and synthetic fragment sequences.

Figure S1. Expression of *CEN1* and *CEN2* in various poplar tissues.

Figure S2. *In vitro* flowering of representative *cen1* mutants.

Figure S3. Phenotypes of soil-grown mutants.

Figure S4. *In vitro* flowering of representative glabrous *cen1cen2myb* (♀) mutants.

Figure S5. Chimeric and abnormal male structure in cen1cen2 (♀) mutants.

Data S1. Summary of mutation patterns determined by amplicon sequencing.

Methods

